# First Genomic Insights into an Aeshnidae Dragonfly: Unveiling the Genome of a Holarctic Species, *Aeshna juncea*

**DOI:** 10.1101/2025.07.15.664918

**Authors:** Ethan Tolman, Dean Bobo, Paul B. Frandsen, Göran Sahlén, Melissa Toups, Jessica L. Ware, Manpreet Kohli

## Abstract

Temperature oscillations in the Arctic may present a unique opportunity to study how insect species respond to such changes. *Aeshna juncea*, a Holarctic species of the family Aeshnidae thrives in this environment; molecular adaptations that allow it to survive in the Arctic have yet to be evaluated. Here, we present the first assembled and annotated draft genome assembly and annotation of *A. juncea*. The assembly is both highly contiguous and complete. This resource is presented and used here to provide further evidence that transposons and unclassified repetitive elements are a major driver behind genome size variation in Odonata and show that the effective population size of *A. juncea* populations from Alaska went through bottlenecks during the most recent ice age. We believe this genome will be an important resource in understanding how species like *Aeshna juncea* survive in Arctic habitats.

## Introduction

The Arctic is a unique environment for insects, with temperatures that frequently swing between cold and warm extremes. Surviving in this environment necessitates both molecular (Danks, 2004; Noer et al., 2023) and behavioral adaptations (Danks, 2004). Arctic insects have long been suggested to be useful indicators of environmental change (Danks, 1992), and many species are currently thought to be in decline (Klimaszewski et al., 2021). Among insects, Odonata, dragonflies and damselflies, are a particularly effective indicator group, and models for evolutionary and ecological study (Córdoba-Aguilar et al., 2023).

*Aeshna juncea* (Linnaeus, 1758) is one such species, with a circumpolar range that spreads from the Arctic southward across Canada, to parts of the United States, Northern Europe and Eurasia. However, the range of *A. juncea* is hypothesized to have shifted in response to climate change (Hickling et al., 2005). Analysis of the Cytochrome Oxidase 1 (COI) haplotypes suggested strong population structure across the Holarctic and divergence times estimated to be around the last glacial cycle, roughly 800,000 years before present (Kohli, Djernæs, et al., 2021). Notably, unique haplotypes endemic to Japan and China were also identified (Kohli, Djernæs, et al., 2021).

*A. juncea* is a dragonfly (Anisoptera) in the family Aeshnidae, which are commonly called darners or hawkers dragonflies. Within Aeshnidae, the genus *Aeshna* consists of ∼30 species: *Aeshna* are large, blue to bluish green dragonflies (Fig. 1a). At its northern range *A. juncea* lives in Arctic bogs in North America and Europe (Fig. 1b) but tends to occupy a variety of habitats when inhabiting the southern regions of these continents (Dijkstra & Schröter, 2020; Wildermuth, 1992) like small ponds with running rivulets and marshes that have stands of sedges (R. Cannings et al., 2000; S. Cannings & Cannings, 1997). *A. juncea* is strongly associated with sedges, hence its common name “Sedge darner”. This species takes at least two years or more to complete its life cycle (Corbet, 1999; Van Buskirk, 1993), but there are likely high rates of plasticity and local variation in response to food availability and temperature (Norling, 1971, 1984). The final instar larvae, that are dominant predators in aquatic habitats, can be large in size but size may vary among populations from 40-51 mm in size (Balázs et al., 2025; Fig 1c. Picture of an *A. juncea* larvae).

**Figure 1:**
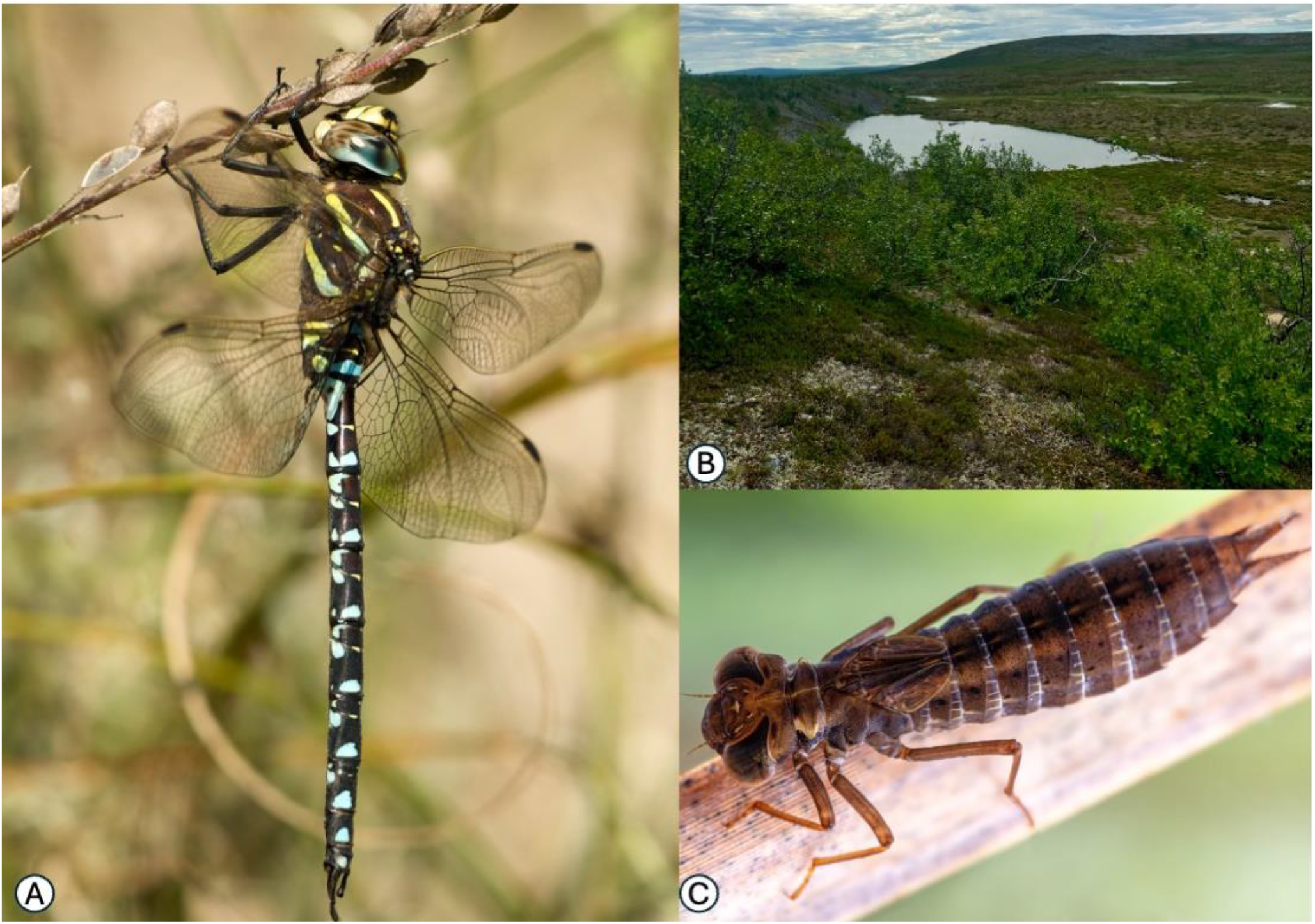
An *A. juncea* adult male (Panel A, note: pictured here is not the individual used in genome sequencing. Picture By Jörg Hempel, CC BY-SA 2.0 de, https://commons.wikimedia.org/w/index.php?curid=4650830), Arctic habitat of *A. juncea* (Panel B, Picture by MKK) and an *A. juncea* larvae (panel C: Photo: Krister Hall).

Presently, there are no published genomes for Arctic Odonata, and no members of the dragonfly family Aeshnidae have a reference genome (NCBI genome search 18th March 2025; Newton et al., 2023). This creates a significant gap in our understanding of molecular patterns in Odonata as Aeshnidae, along with Austropetalidae, are the earliest branching extant lineage within Anisoptera (Bybee et al., 2021; Goodman et al., 2023; Kohli, Letsch, et al., 2021; Suvorov et al., 2021). Filling in these gaps is critical work, as high-quality reference genome assemblies are a fundamental tool in understanding organismal evolution (Newton et al., 2023). Despite the limited number of published Odonata genomes (20 genomes: 9 Anisoptera and 11 Zygoptera, NCBI - genome database accessed March 18th 2025), important insights into Odonata ecology and evolution have already emerged, including: the discovery of highly conserved chromosome structure across suborders (Tolman, et al., 2023), the detection of unexpected hybrid zones (Patterson et al., 2024), and the identification of expanded gene families implicated in migratory behavior and persistence in urban habitats (Tolman et al., 2025). This first Aeshnidae genome is not only helpful for new insights into odonate evolution but also takes us a step forward in understanding species inhabiting the Arctic.

Here we present a high-quality draft genome assembly of *A. juncea*, use the genome to model the historical demography of *A. juncea*, and consider some of the implications of the genome in the study of the evolution of Odonata.

## Methods

### Specimen collection and Sequencing

An *A. juncea* adult was collected in Fairbanks, Alaska (64.819627, -147.869695) on 17th July 2023. The adult female specimen was flash-frozen in liquid nitrogen and transported to the BYU DNA sequencing center. The sequencing center extracted high molecular weight DNA with a Qiagen Genomic Tip DNA extraction kit. Purified DNA was sheared to 18 kbp using a Diagenode Megaruptor and fractions >10kbp were selected using a SAGE Science Blue Pippin. The sequencing library was then prepared using the PacBio HiFi SMRTbell Express Template Kit 2.0. The library was sequenced on a single PacBio Revio SMRT cell.

### Genome assembly and QC

We estimated the genome assembly with Genomescope 2.0 (Ranallo-Benavidez et al., 2020) after generating read histograms with Jellyfish v2.2.10 (Ahrens et al., 2018; Marçais & Kingsford, 2011) with a Kmer size of 21. We assembled the HiFi reads into a draft assembly using hifiasm v.0.16.0 using default flags (Cheng et al., 2021). Contaminants were identified and removed as a part of the NCBI online contamination screen (supplementary table 1). We generated all contiguity stats for the contaminant-free assembly with assembly-stats v0.1.4 (*Assembly-Stats*, 2014/2022). Completeness of the cleaned assembly was assessed through BUSCO v.4.1.4 (Manni et al., 2021) using the Insecta ODB10 database (Kriventseva et al., 2019. We ran BUSCO in genome mode with the flag **-**–long, to train Augustus using the initially identified orthologues, in order to more accurately identify all sets of core orthologues. We calculated coverage by mapping our reads to the cleaned assembly with minimap2 v2.1 (Li, 2018) using default flags.

### Genome Annotation

Repetitive elements in the genome assembly were modeled and soft-masked with RepeatModeler2 v2.0.1 and RepeatMasker v.4.1.2-p1 (Flynn et al., 2019). We then identified protein coding sequences with the program GALBA (Brůna et al., 2023) using the annotated protein set of *Pantala flavescens*, the most high quality annotation for an Odonata genome to date, as a reference (H. Liu et al., 2022). We then discarded annotated genes that did not have a significant BLAST (Camacho et al., 2009) hit (with an e-value cutoff of 1e-10) to a protein in the reference set, which reduces annotation specificity with only a slight decrease in sensitivity (Tolman et al., 2024). The justification for this annotation approach, including the use of GALBA over approaches that leverage transcriptomic data, the use of *P. flavescens* as a reference for Odonata annotations, and discarding annotated proteins that are not found in the reference set are more thoroughly discussed in Tolman et al. (2025). We also identified the mitochondrial genome using mitohifi v. 3.0.0 (Uliano-Silva et al., 2021), using the mitochondrial genome of the aeshnid dragonfly *Anax imperator* (Herzog et al., 2016) as reference.

### Analysis of Demographic History

To consider how *A. juncea* may have responded to past changes in climate, we modeled the change in historical effective population size of *A. juncea* (N_e_) using the Pairwise Sequential Markovian Coalescent (PSMC). We utilized a pipeline and parameters that have previously been used to model historical demography for Odonata sequenced using PacBio HiFi technology with the PSMC (Tolman, Bruchim, et al., 2023; Tolman et al., 2024). Briefly, we converted the previously generated bam file (generated by mapping the reads to the genome assembly) into a sam file with samtools v1.16.1 (Danecek et al., 2021). We called bases with a minimum depth of 15 and a maximum depth of 100 with samtools v1.16.1 and bcftools v1.6 (Danecek et al., 2021) using default filtering settings to avoid possible contaminants. We then converted the basecalling output to psmcfa format using psmc v0.6.5-r67 (S. Liu & Hansen, 2017) and then ran the PSMC with 100 bootstrap replicates, splitting the first time window into two windows to avoid false positive peaks (Hilgers et al., 2025). We visualized the resulting demographic with a gnu-plot v5.2 (Phillips, 2012) using a mutation rate of 2e-9 (an approximately average rate for insects (Liu et al., 2017)), and a generation time of 2.5 years (Van Buskirk, 1993).

## Results

### Genome Assembly and QC

We obtained > 6 million reads for a total of >75 Gb (N50=12944, L50=2331820, ave=12379.94, 6094406). The estimated genome size from Kmer coverage was ∼1.46 Gb (Supplementary Fig. 1). The NCBI contamination screen removed 24 contigs identified as contaminants, all assigned to firmicutes, totaling over 2.7 Mb. The blobplot analysis showed evidence of contamination from Pseudomonadota, with such contigs showing both low coverage and GC content (Supplementary Fig. 2). Contaminants assigned to Pseudomonadota were removed from the genome assembly. The resulting clean assembly was just over 1.9Gb in size, highly contiguous (N50>16Mb, L50=27, n=1571, ave>1.2 Mb), and contained 98% of insecta ODB10 orthologues, with low levels of duplication (C:98.0%[S:95.2%,D:2.8%],F:0.7%,M:1.3%,n:1367). The genome assembly averaged 38x read coverage.

### Genome Annotation

Repetitive elements made up 46.43% of the genome assembly, with unclassified repetitive elements (26.78% of the assembly), Retroelements (12.16% of the assembly), and DNA transposons (5.97% of the assembly) making up the bulk of the repetitive elements (Table 1). These repetitive elements were largely composed of very recently (Kimura substitution level < 10) and moderately recently (Kimura substitution level 10< >30) diversifying families of repetitive elements (Fig. 2). The filtered annotation set contained 16,675 transcripts, with most of the insecta ODB10 orthologues present (C:98.2%[S:95.2%,D:3.0%],F:0.4%,M:1.4%,n:1367).

**Table One:** Repetitive Elements in the Draft Genome Assembly of *A. juncea*.

| Table One: Repetitive Elements in the Draft Genome Assembly of *A. juncea* |  |  |  |
| --- | --- | --- | --- |
| Category | Number of Elements | Length Occupied | Percentage of genome assembly |
| **Retroelements** | 843945 | 232720612 bp | 12.16% |
| *SINEs* | 24765 | 1962901 bp | 0.10% |
| *Penelope* | 167766 | 28993412 bp | 1.52% |
| *LINES* | 786232 | 218336891 bp | 11.41% |
| CRE/SLACS | 0 | 0 bp | 0.00% |
| L2/CR1/Rex | 312761 | 75575218 bp | 3.95% |
| R1/LOA/Jockey | 25408 | 16846890 bp | 0.88% |
| R2/R4/NeSL | 155 | 50521 bp | 0.00% |
| RTE/Bov-B | 265041 | 86571217 bp | 4.52% |
| L1/CIN4 | 337 | 391387 bp | 0.02% |
| *LTR elements* | 32948 | 12420820 bp | 0.65% |
| BEL/Pao | 738 | 643223 bp | 0.03% |
| Ty1/Copia | 8096 | 5052634 bp | 0.26% |
| Gypsy/DIRS1 | 9240 | 5171092 bp | 0.27% |
| Retroviral | 14874 | 1553871 bp | 0.08% |
| **DNA transposons** | 491044 | 114165405 bp | 5.97% |
| *hobo-Activator* | 213487 | 42985991 bp | 2.25% |
| *Tc1-IS630-Pogo* | 140827 | 35041086 bp | 1.83% |
| *En-Spm* | 0 | 0 bp | 0.00% |
| *MuDR-IS905* | 0 | 0 bp | 0.00% |
| *PiggyBac* | 25484 | 12758092 bp | 0.67% |
| *Tourist/Harbinger* | 5703 | 772016 bp | 0.04% |
| *Other (Mirage, P-element, Transaib)* | 8611 | 2097024 bp | 0.11% |
| **Rolling-circles** | 83604 | 19017078 bp | 0.99% |
| **Unclassified** | 2144714 | 512505102 bp | 26.78% |
| **Total interspersed repeats** | - | 859391119 bp | 44.91% |
| **Small RNA** | 5504 | 3293839 bp | 0.17% |
| **Satellites** | 26129 | 3172044 bp | 0.17% |
| **Simple repeats** | 76307 | 3504080 bp | 0.18% |
| **Low complexity** | 0 | 0 bp | 0.00% |

**Figure 2:**
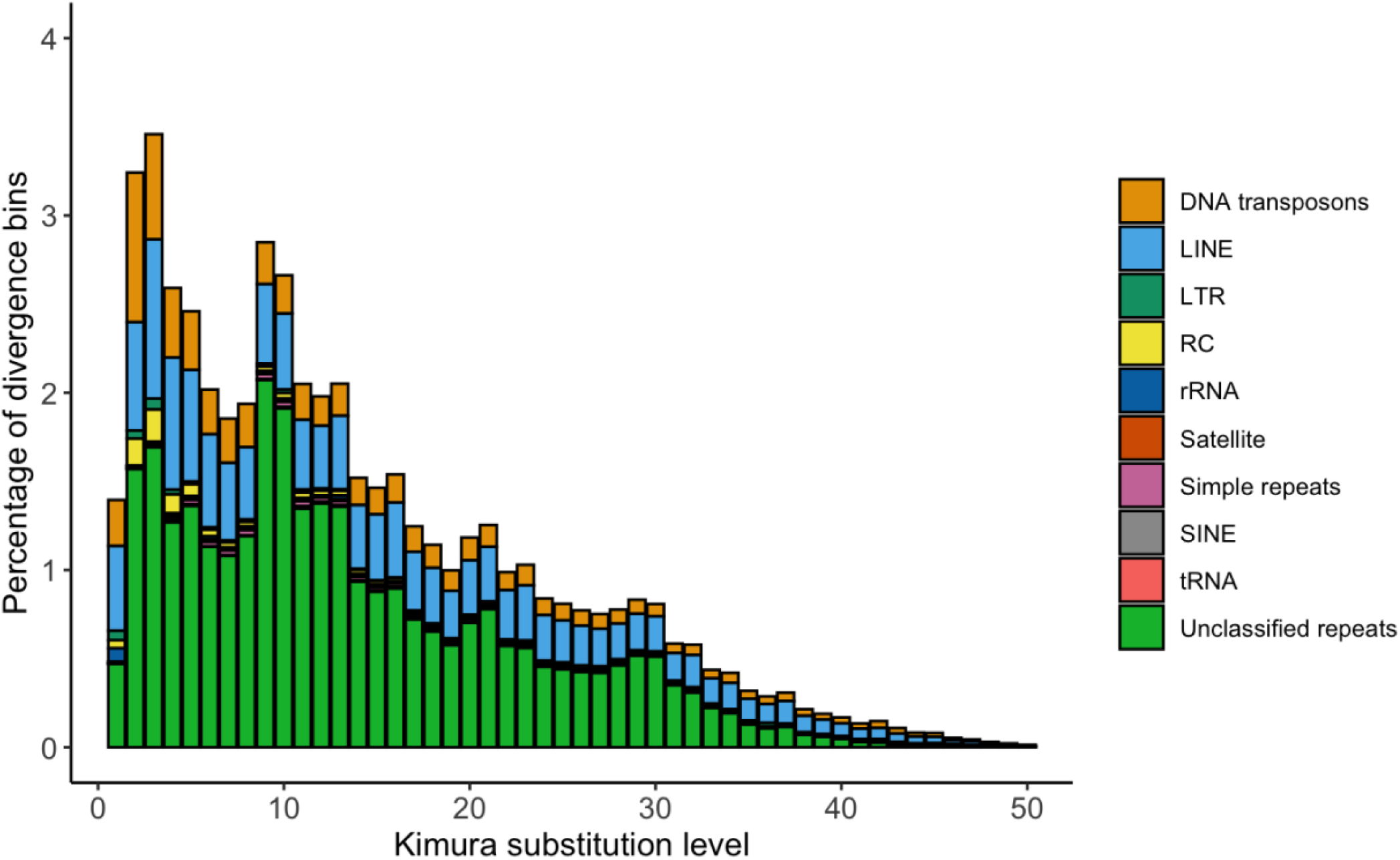
Repeat landscape of A. juncea. Families of repetitive elements are clustered according to Kimura substitution level, and then colored by classification.

The demographic model showed evidence of a series of strong bottlenecks <100kya, resulting in a greater than 90% reduction in effective population size, N_e,_ to ∼ 3,000 (Fig. 3). Prior to this bottleneck, the N_e_ was estimated to range between 400 thousand and 1.2 million (Fig. 2).

**Figure 3:**
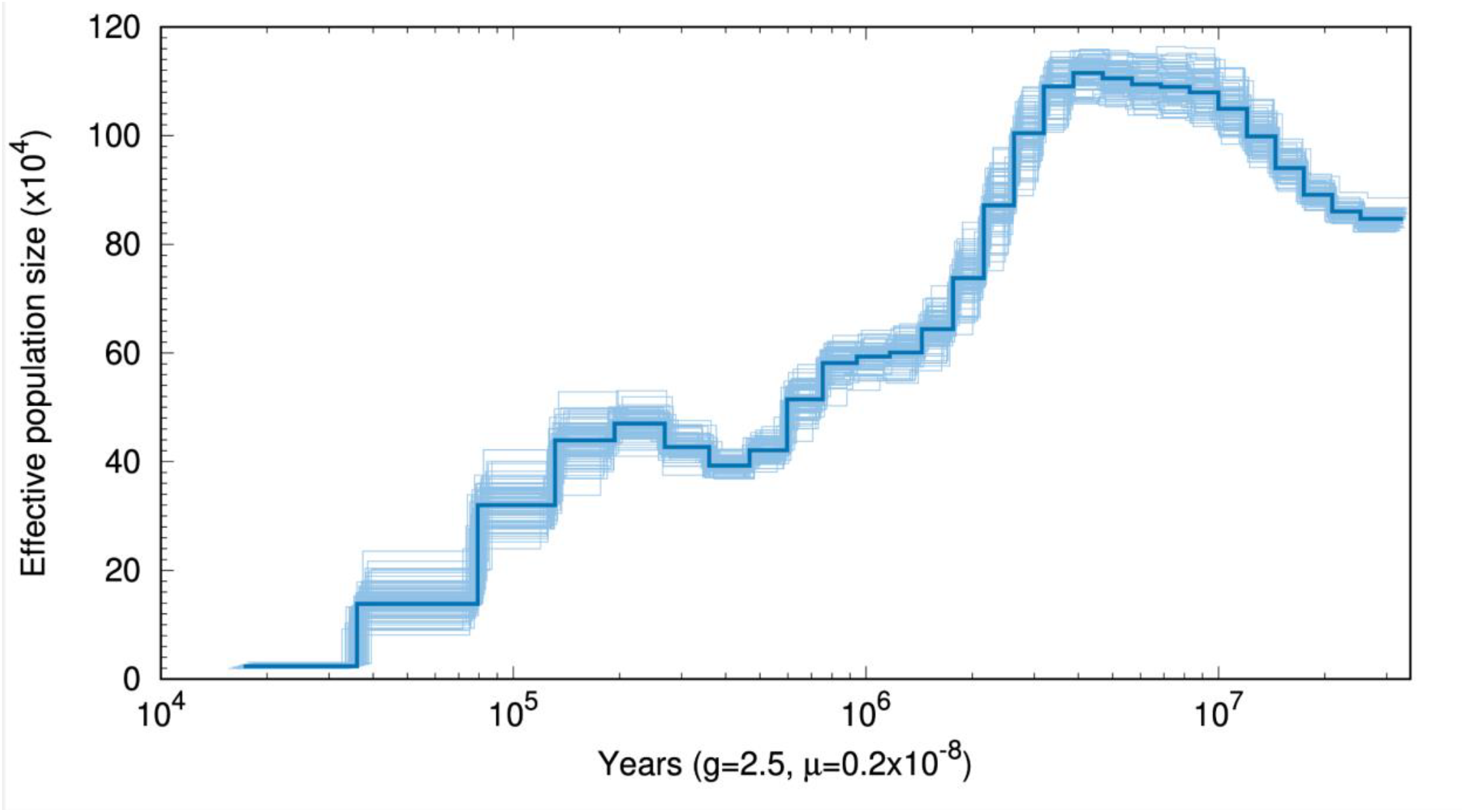
Historical Demography of A. juncea. Changes in N_e_ are shown as modeled with the Pairwise Sequential Markovian Coalescent (Liu & Hansen, 2017).

## Discussion

### The largest Odonata genome assembly to date

At over 1.9 Gb, the genome assembly of *Aeshna juncea* is the largest of any Odonata sequenced to date. Of comparable size are the genome assemblies of the damselflies *Platycnemis pennipes* (1.79 Gb) and *Ischnura elegans* (1.72 Gb) and the dragonfly *Tanypteryx hageni* (1.70 Gb) (Tolman, Beatty, et al., 2023). In each of these three previously sequenced Odonata, expanded repetitive elements (particularly retroelements, DNA transposons and unclassified repeats) are largely responsible for an increased genome size; in these taxa repetitive elements made up more than 40% of each genome assembly, while by contrast repetitive elements comprised less than 20% of the relatively small genome assembly of *P. flavescens* (0.67 Gb; Tolman et al., 2023). Total interspersed repetitive elements make up 44.91% of the genome assembly of *Aeshna juncea* (Table 1), adding support to the hypothesis that repetitive elements are the primary engine of genome size expansion in Odonata, as is the case in other studied insect orders (Heckenhauer et al., 2022; Yuan et al., 2024; Zhao et al., 2025). The implications of this hypothesis in relation to the species, ecological, morphological and behavioral diversity of Odonata cannot yet be fully explored given the limited number of Odonata genome assemblies. However, a study of the role of repetitive elements in the biodiversity of Odonata would be an exciting next step in Odonatology, and in entomology more generally (Sproul et al., 2023).

### Historical demography of Aeshna Juncea

The N_e_ of *A. juncea* showed a series of dramatic reductions from 100-20 Ka (roughly overlapping with the last ice age), well after the populations from different geographic regions are estimated to have diverged (Kohli, Djernæs, et al., 2021). However, we caution that more recent coalescent events cannot be extended to other populations of *A. juncea*, while deeper coalescent events are expected to converge with other *Aeshna* sharing a MRCA with *A. juncea* less than 20 Ma. It is important to note that this model is based upon estimates of genome-wide mutation rates that have not been empirically tested in Odonata. Different rates change our estimates of the timing of bottleneck events, but they do not typically erase coalescent events, and bias is found to be less pronounced in the more recent past (Tolman, Bruchim, et al., 2023). Identifying the drivers behind this collapse in N_e_ is important for understanding the barriers between populations, and how *A. juncea* may fare on a rapidly changing planet. It is unclear if the Arctic populations are currently dropping in N_e_ due to warming, but this finding does warrant further investigation.

### Future directions in Arctic Odonata

This genome assembly is an important resource in studying the evolution of Arctic Odonata. Population level resequencing data from across the species range has the potential to identify loci under selection in Arctic populations implicated in survival of this hostile habitat. A comparison of this genome assembly with the genome assemblies of exclusively temperate *Aeshna* will be an important exercise to identify expanded gene and repeat families that could be implicated in Arctic survival. Most importantly, repeating these experiments with other Arctic species (and determining if other Arctic species and populations exhibit the same contraction in N_e_ around the Chelford Interstadial) will provide a strong foundation for critical research into Arctic species.

## Data availability

The genome assembly and reads are available on NCBI (bioproject: PRJNA1156168), while the genome annotations have been uploaded to FigShare (10.6084/m9.figshare.29360816).

## Acknowledgments

This work was supported by funds from Baruch College and PSC-CUNY-Cycle54 grant to Kohli. We also thank Dr. Anton Suvorov for supporting this project.

