## Supplementary Materials for "First Genomic Insights into an Aeshnidae Dragonfly: Unveiling the Genome of a Holarctic Species, *Aeshna juncea*"

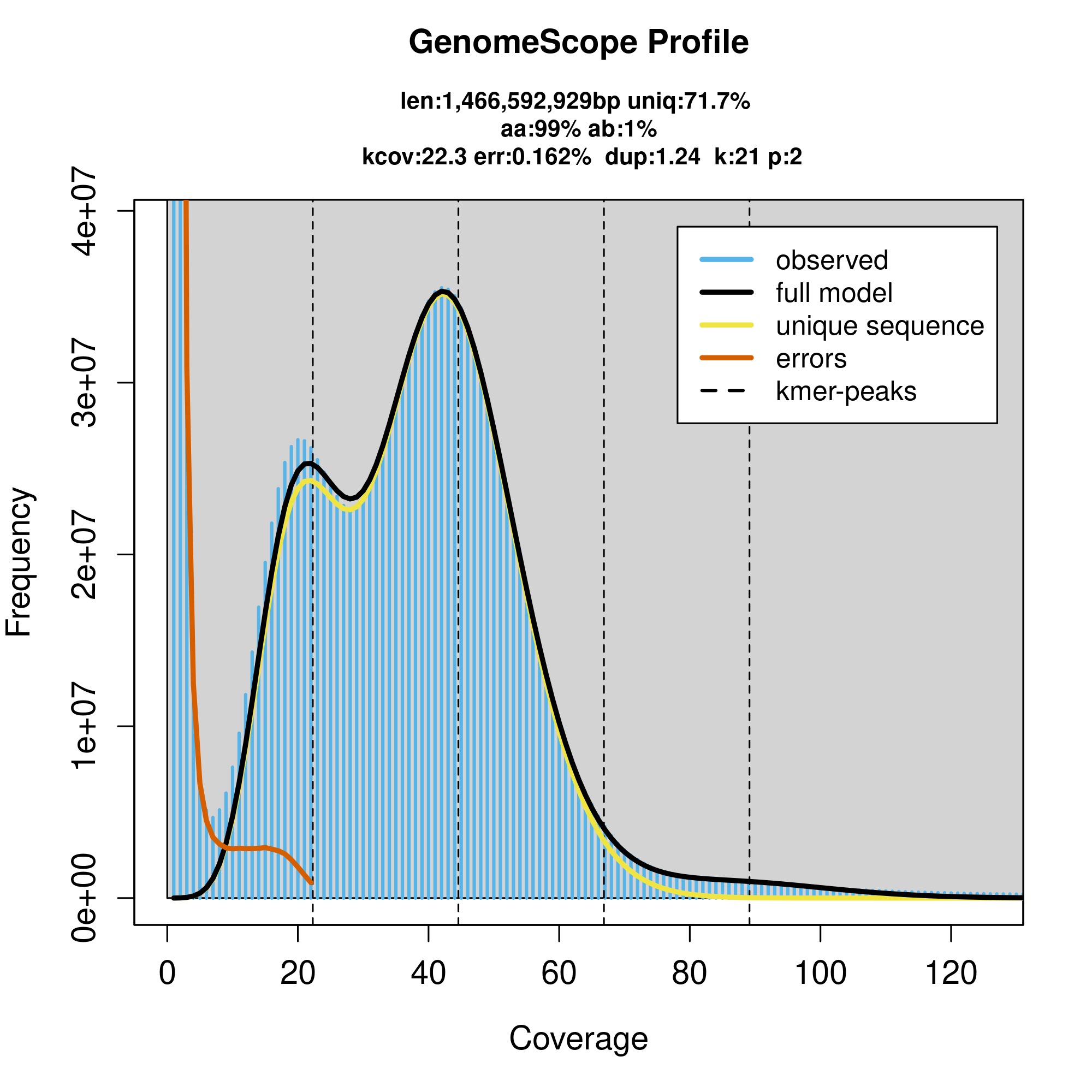

*Supplementary Figure 1: Genome Scope profile of PacBio Hifi reads generated from A. juncea. Heterozygous and homozygous peaks are shown at ~19x and 38x coverage respectively.*

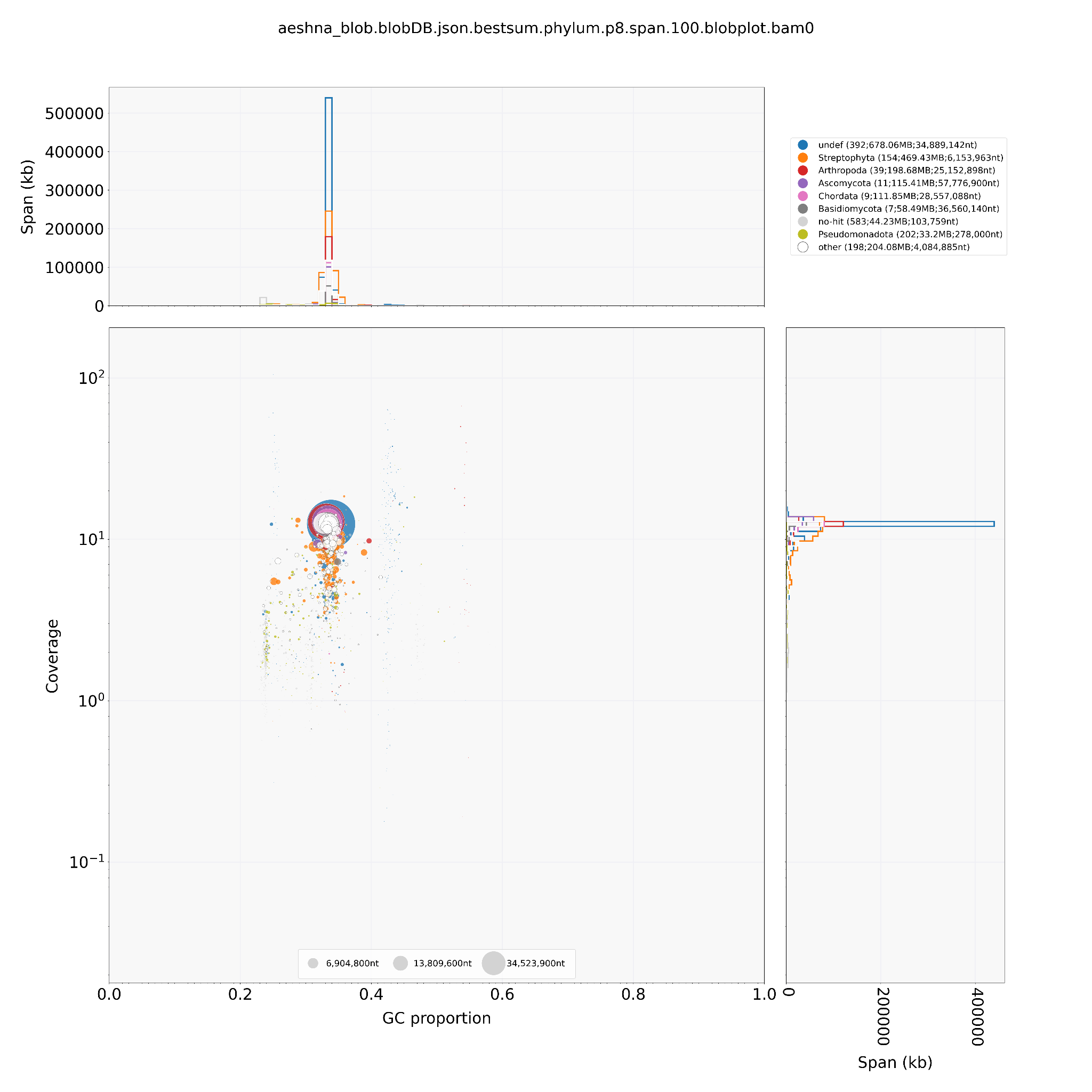

*Supplementary Figure 2: Taxon-annotated coverage plot of A. juncea genome.*

| Supplementary Table 1: Contaminants Identified by the NCBI Contamination Screen | | |
| --- | --- | --- |
| Contig | Contig length | Contamination source |
| ptg000235l | 169541 | prok:firmicutes |
| ptg000304l | 83800 | prok:firmicutes |
| ptg000397l | 651572 | prok:firmicutes |
| ptg000419l | 185203 | prok:firmicutes |
| ptg000425l | 54706 | prok:firmicutes |
| ptg000434l | 198457 | prok:firmicutes |
| ptg000517l | 144871 | prok:firmicutes |
| ptg000526l | 128014 | prok:firmicutes |
| ptg000536l | 212888 | prok:firmicutes |
| ptg000578l | 85950 | prok:firmicutes |
| ptg000667l | 102334 | prok:firmicutes |
| ptg000691l | 68758 | prok:firmicutes |
| ptg000726l | 90641 | prok:firmicutes |
| ptg000900l | 98213 | prok:firmicutes |
| ptg000951l | 77181 | prok:firmicutes |
| ptg000976l | 29441 | prok:firmicutes |
| ptg000988l | 106979 | prok:firmicutes |
| ptg001018l | 44302 | prok:firmicutes |
| ptg001260l | 32196 | prok:firmicutes |
| ptg001263l | 21774 | prok:firmicutes |
| ptg001429l | 74166 | prok:firmicutes |
| ptg001463l | 57991 | prok:firmicutes |
| ptg001492l | 20429 | prok:firmicutes |
| ptg001565l | 29028 | prok:firmicutes |
| ptg000163l | 1308677 | Adaptor sequence |
| ptg000264l | 2287487 | Adaptor sequence |
